# Codependency and mutual exclusivity for gene community detection from sparse single-cell transcriptome data

**DOI:** 10.1101/2021.03.15.435370

**Authors:** Natsu Nakajima, Tomoatsu Hayashi, Katsunori Fujiki, Katsuhiko Shirahige, Tetsu Akiyama, Tatsuya Akutsu, Ryuichiro Nakato

## Abstract

Single-cell RNA-seq (scRNA-seq) can be used to characterize cellular heterogeneity in thousands of cells. The reconstruction of a gene network based on coexpression patterns is a fundamental task in scRNA-seq analyses, and the mutual exclusivity of gene expression can be critical for understanding such heterogeneity. Here, we propose an approach for detecting communities from a genetic network constructed on the basis of coexpression properties. The community-based comparison of multiple coexpression networks enables the identification of functionally related gene clusters that cannot be fully captured through differential gene expression-based analysis. We also developed a novel metric referred to as the exclusively expressed index (EEI) that identifies mutually exclusive gene pairs from sparse scRNA-seq data. EEI quantifies and ranks the exclusive expression levels of all gene pairs from binary expression patterns while maintaining robustness against a low sequencing depth. We applied our methods to glioblastoma scRNA-seq data and found that gene communities were partially conserved after serum stimulation despite a considerable number of differentially expressed genes. We also demonstrate that the identification of mutually exclusive gene sets with EEI can improve the sensitivity of capturing cellular heterogeneity. Our methods complement existing approaches and provide new biological insights, even for a large, sparse dataset, in the single-cell analysis field.

## 1 Introduction

Single-cell RNA sequencing (scRNA-seq) enables us to explore and characterize variability in individual cells. At the same time, it can provide information on the regulatory relationships between genes and how genes interact with each other at the single-cell level. Since scRNA-seq experiments profile the dynamics and variation of gene expression in different cell states, such as the cell cycle, cell division and cell differentiation, across thousands of cells, the reconstruction of gene regulatory networks (GRNs) helps us to understand gene functions and cell type- or state-specific variability in genetic interactions.

Many mathematical methods have been proposed to infer GRNs from bulk transcriptome data, including the use of Boolean networks [1, 2, 3, 4], Bayesian networks [5, 6, 7, 8], mutual information [9, 10, 11] and linear regression [12, 13]. Although most of these methods enable us to capture codependency and regulatory interactions from a dataset with a limited sample size, they are not suitable for inferring regulatory relationships on the basis of temporal information or sparse expression. In addition, they are not applicable to large-scale networks due to their high computational complexity.

With the development of high-throughput sequencing techniques, expression can now be measured simultaneously in a large number of cells. In particular, while droplet-based single-cell RNA sequencing can measure expression levels in thousands of cells, it exhibits lower sensitivity for each gene compared to other scRNA-seq methods [14]. This leads to an excessive amount of zero read counts, which is likely to affect downstream analysis. Previous studies as mentioned in [15, 16] are useful for integrated datasets generated by distinct platforms. The lower sensitivity of sparse scRNA-seq might be improved by integrating datasets from several distinct platforms. However, it would be better to develop methods to capture the specific expression from sparse scRNA-seq data through a single platform.

For the inference of GRNs from scRNA-seq data, numerous methods have been developed [17, 18, 19, 20, 21, 22]. These methods are mostly aimed at reconstructing the statistical dependency between genes based on the ordering of single cells according to the time information underlying dynamic processes from a dataset obtained from high-quality cells. The variability in gene expression can be revealed by taking advantage of underlying temporal information for individual cells. In addition, an approach based on information theory estimates non-linear dependencies with pairwise joint probability distributions if the sample size is sufficient [17]. However, the sparsity of single-cell data might mean that these data are not inherently suitable for the inference of GRNs, and GRN algorithms are mostly effective for up to a thousand genes [23].

Moreover, the community detection of coexpression networks is important for the identification of groups of functionally related genes. To address the community detection problem, many methods have been studied [24, 25, 26, 27, 28]. The Girvan-Newman method basically decomposes a network to maximize the modularity [24, 29]. To improve the computational complexity, the Leading eigenvector method, which partitions a network based on the spectral optimization of modularity according to the eigenvector, is developed [27].

Similar to codependency, mutual exclusivity is an inherent expression pattern in gene expression. At single-cell resolution, mutually exclusive genetic alterations are likely to be one cause of cellular heterogeneity not only within a tumour but also during cell development. Notably, several approaches have been developed for detecting mutually exclusive gene sets associated with cancer driver mutations. For example, mutual exclusivity modules (MEMo) are used to identify gene sets that belong to the same pathway by extracting all groups of functionally related genes from the Human Reference Network using multiple somatic mutation datasets [30]. Other methods predict whether the cause of an amino acid change is associated (or not) with cancer based on a random forest with somatic missense mutations [31, 32]. Although these methods are suitable for detecting independent alterations in cancer driver genes from mutation data, the modelling of mutual exclusivity from scRNA-seq data has yet to be studied.

In this paper, we focus on the modelling of one-to-one relationships between two genes, such as codependency and mutual exclusivity, because codependency and mutual exclusivity are inherent and specific gene expression patterns. The purpose of this paper is twofold. First, we develop an approach to detect communities of coexpression networks with the co-dependency index (CDI) [33]. We evaluate the effectiveness of community detection based on glioblastoma scRNA-seq data and show that this approach finds not only densely connected subgraphs but also functionally related networks. Communitybased comparisons provide information on the similarities and differences in coexpressed genes when applied to multiple samples. Second, we develop a novel metric, the exclusively expressed index (EEI), for identifying mutually exclusive gene sets from sparse scRNA-seq data. According to the idea of detecting sparse expression [33], EEI enables us to quantify mutually exclusive expression due to negative correlations and genetic alterations not only in cancer driver genes but also in cell-type specific marker genes if the input is scRNA-seq data. We apply our method to glioblastoma scRNA-seq data and show that it outperforms existing methods while maintaining robustness against the sequencing read depth, and mutually exclusive gene sets can improve the sensitivity of the identification of cellular heterogeneity.

## 2 Methods

### 2.1 Co-Dependency Index Score

A previous study proposed a method, the co-dependency index (CDI), for quantifying the codependency relationship between two genes, gene *i* and gene *j*, across thousands of cells from scRNA-seq data [33].

Assuming that *g_i_* and *g_j_* are independently expressed, *P*(*g_i_* = 1) denotes the probability of observing that *g_i_* has the nonzero expression. *P*(*g_i_* = *1,g_j_* = 1) denotes the joint probability of observing that both genes *g_i_* and *g_j_* exhibit coincident nonzero expression values in the same cell and is formulated as follows:

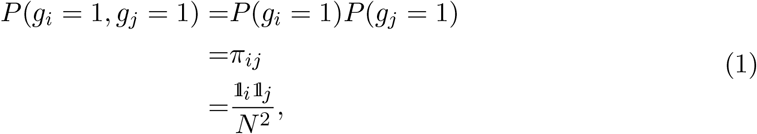

where 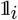 and *N* represent the number of cells in which *g_i_* presents nonzero values and the total number of cells, respectively. Under this assumption, *p_e_*(*π_ij_*), defined as follows, is the probability of observing a test statistic being as extreme under the null hypothesis that genes *g_i_* and *g_j_* are independent:

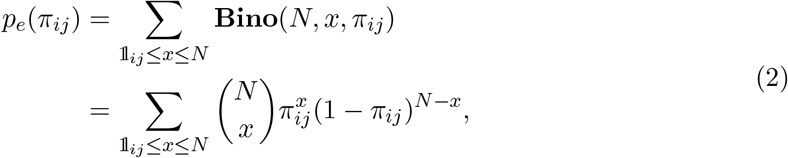

where **Bino**(*N, x, π_ij_*) represents the probability of observing *x* successes in *N* trials if the probability of success is *π_ij_*. 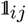 is the number of cells in which *g_i_* and *g_j_* simultaneously present nonzero values. Then, CDI is defined as follows:

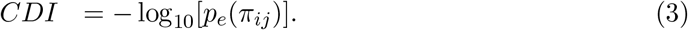

### 2.2 Exclusively Expressed Index Score

#### 2.2.1 Definition

In this study, we propose a novel metric, the exclusively expressed index (EEI), for the quantification of mutual exclusivity between two genes according to the concept of CDI from sparse scRNA-seq data. Mutually exclusive expression between two specific genes, *g_i_* and *g_j_*, can be divided into two cases: A and B. In the case of A, *g_i_* shows zero expression, while *g_j_* shows nonzero expression, and the probability that A will occur can be denoted by *P*(*g_i_* = *0,g_j_* = 1). In the case of B, *g_i_* exhibits nonzero expression, *g_j_* exhibits zero expression, and *P*(*g_i_* = 1,g_j_ = 0) indicates the probability of B. EEI is computed for each pair of genes for all possible combinations. Under the null hypothesis that *g_i_* and *g_j_* are independent, the probability of the occurrence of A is defined as follows:

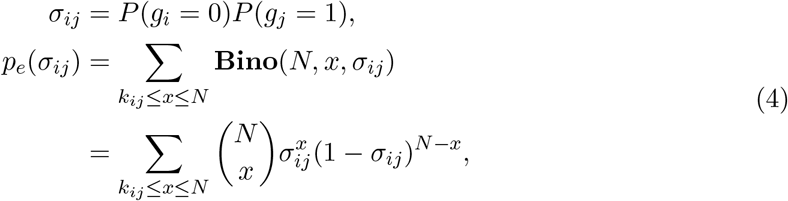

where *k_ij_* represents the number of cells in which *g_i_* presents zero values and *g_j_* presents nonzero values. Similarly,

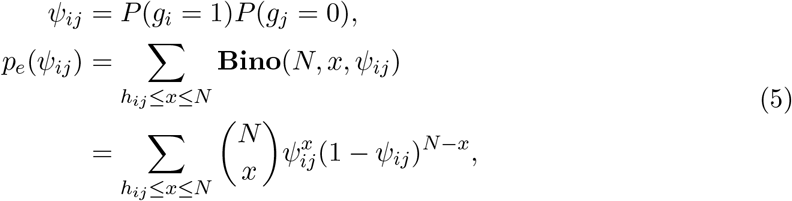

where *h_ij_* represents the number of cells in which *g_i_* presents nonzero values and *g_j_* presents zero values. Then, EEI is defined as follows:

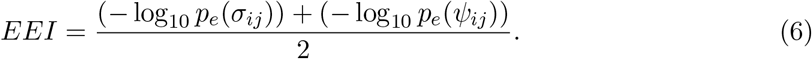

Although CDI and EEI do not impute technical zeros, these metrics calculate the possibility of codependency and mutual exclusivity by counting the number of cells according to binary quantification if the sample size is sufficient. Gene pairs with lower CDI scores cannot be applied to alternative gene pairs with higher EEI scores. Since CDI captures the possibility of coexpression rather than being a coincidence, the gene pairs that are expressed in all cells (i.e., housekeeping genes) exhibit lower CDI and EEI scores. A high EEI can directly quantify that the gene pairs exhibit mutually exclusive expression.

#### 2.2.2 Application of EEI to Single-Cell Clustering

Mutually exclusive expression can be observed as genetic alterations [34, 35, 36, 37] and negative correlations [38, 39] between two specific genes. Genetic alterations lead to protein production because mutually exclusive gene pairs might be amplified in different cell populations [34]. Notably, tumour cell heterogeneity might be considered to be caused by mutations in driver genes or to result from tumour cell progression. At single-cell resolution, mutually exclusive expression means that it is highly possible that two genes are exclusively expressed in the different cell types. Since this property is specific to singlecell populations, we attempt to apply the expression of mutually exclusive gene sets to improve the sensitivity of the identification of cellular heterogeneity. In this subsection, we introduce a novel strategy for the classification of single cells with EEI gene sets.

Clustering analysis with EEI can be summarized as follows.

1. EEI is calculated for all gene pairs from scRNA-seq data, and the 1,000 gene sets with the highest EEI scores are extracted.
2. To generate a feature matrix, the expression ratio (i.e., the proportion of the expression values) is calculated using normalized (i.e., normalize the gene expression for each cell by the total read count) and log-transformed expression for all extracted pairs. Then, the features of expression are merged with the log-transformed expression values in the scRNA-seq dataset, as shown in Figure 1A.
3. The merged data can be regarded as an input, and classification is performed via dimensionality reduction methods. In this experiment, we reduce the high-dimensional data to 40 dimensions via singular value decomposition (SVD), and then UMAP is used to reduce 40 dimensions to 2 dimensions.

**Figure 1:**
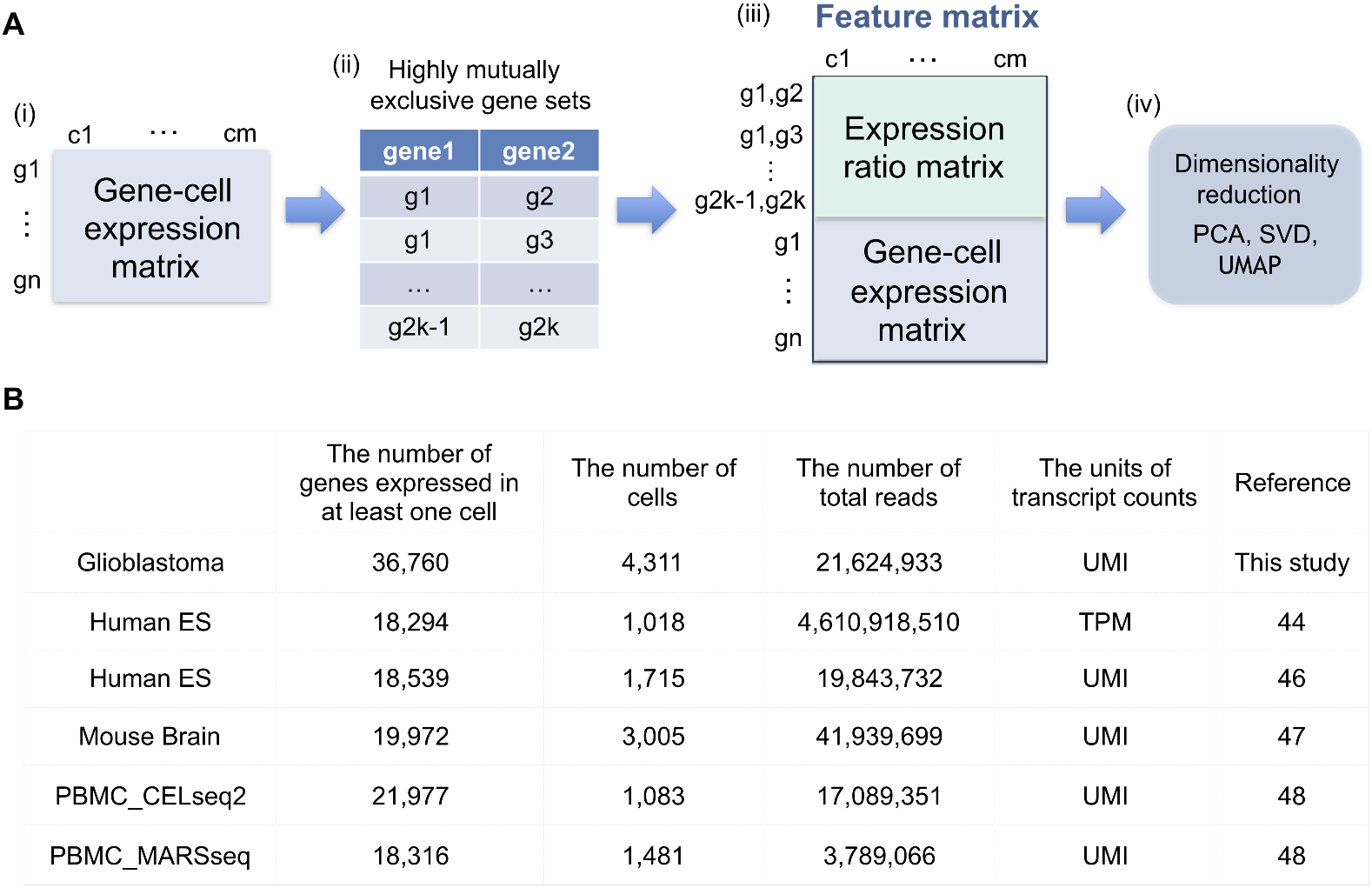
(**A**) Overview of the generation of a feature matrix from scRNA-seq data. (i)(ii) If a gene-cell expression matrix is provided, EEI is calculated, and highly mutually exclusive gene pairs are extracted. (iii) The feature matrix is generated by merging the expression ratio matrix for EEI pairs with the gene-cell expression matrix. (iv) Dimension reduction is performed using SVD and UMAP with the feature matrix as an input. (**B**) Summary of the six scRNA-seq datasets. This contains the number of genes that expressed in at least one cell, the number of cells, the number of total reads, the units of transcript counts and the reference.

### 2.3 Community Detection Algorithm

Community detection can extract topological characteristics from complex networks. The Girvan-Newman method is a basic method that decomposes a network iteratively by removing the edges connecting communities that present the highest edge betweenness [24, 29] based on maximizing modularity, Q [24]:

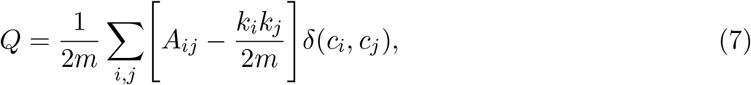

where *m* represents the total number of edges, *A_ij_* is the weight of the edge between nodes *i* and *j*, *k_i_* is the sum of the weights of edges attached to node *i* or the degree of node *i*, *c_i_* is the community to which node *i* belongs, and *δ* is defined as *δ*(*u,v*) = 1 if *u* = *v* and as 0 otherwise.

The Leading eigenvector method [27] is based on the spectral optimization of modularity by reformulation of modularity in terms of the eigenvalue [40]. It calculates the eigenvector of the modularity matrix and the largest positive eigenvector partitions the network into two communities. This algorithm performs faster than other modularity optimizations and slightly better for large-scale networks and a wide variety of networks [41, 42].

This method maximizes modularity, which is defined in terms of a matrix based on the eigenvalues and eigenvectors, called the modularity matrix, B [29]:

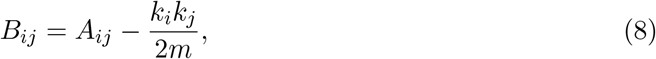

where *A* is the adjacency matrix. Then, modularity is defined as follows:

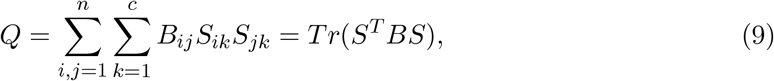

where *S* is an index matrix to detect *c* (*c* ≥ 2) communities. Each column of this matrix is an index vector of (0,1) elements. Writing *B* = *U DU^T^*, where *U* = (*u*_1_|*u*_2_|…) is the matrix of eigenvectors of *B, D* is the diagonal matrix of eigenvalues *D_ii_* = *β_i_* and *s* is an index vector of (−1,1) elements in which 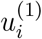 is the i-th element of the eigenvector, *u_1_* of *B*, then *Q* is defined as follows:

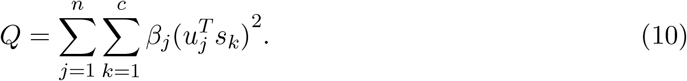

For a different approach, the Louvain method [29] is an agglomerative hierarchical clustering method to maximize modularity by local optimization. It aggregates each cluster into a single node until the modularity does not increase, which leads to a small-sized network and fast performance [43].

In this study, we applied the Leading eigenvector method because it is a fundamental method based on spectral optimization that partitions a network into clusters with eigenvalue decomposition.

### 2.4 Dataset

To evaluate the performance of EEI and the community detection of coexpression networks, we applied six scRNA-seq datasets.

Glioblastoma scRNA-seq data: We generated single-cell RNA-seq data obtained from glioblastoma stem-like cells before (stem) and after (serum^+^) the addition of serum that are collected at 0 and 12 hours (see Section 2.5). Glioblastoma stem-like cells are subsets of glioblastoma cells that possess self-renewal ability and exhibit extensive tumorigenicity. The datasets obtained at 0h and 12h contained 2,102 and 2,209 single cells in total, respectively.

Human ES progenitor scRNA-seq data: This dataset was published by Chu *et al.* [44] and provides snapshots of lineage-specific progenitor cells differentiated from human ES cells. The progenitor cells consisted of 1,018 single cells in total and included cell types such as neural progenitor cells, endoderm cells, endothelial cells and trophoblast-like cells. Library preparation was performed using the Fluidigm C1 system [45].

Human ESC-derived neuron scRNA-seq data: This dataset was published by Manno *et al.* [46] and provides the transcriptomes of human ventral midbrain single cells. These cells, 1,715 in total, included cell types such as oculomotor and trochlear nucleus, serotonergic and medial neuroblasts.

Mouse cortex scRNA-seq data: This dataset was published by Zeisel *et al.* [47] and provides the transcriptomes of mouse cortex and and hippocampal cells. These cells, 3,005 in total, included cell types such as interneurons, oligodendrocytes and microglial cells.

PBMC CELseq2 scRNA-seq data: This dataset was published by Mereu *et al.* [48] and provides a reference sample containing human peripheral blood mononuclear cells that were generated with the CELseq2 protocol. These cells, 1,083 in total, included cell types such as B cells, NK cells and CD14 monocytes.

PBMC MARSseq scRNA-seq data: This dataset was published by Mereu *et al.* [48] and provides a reference sample containing human peripheral blood mononuclear cells that were generated with the MARSseq protocol. These cells, 1,481 in total, included cell types such as CD4 T cells, FCGR3A monocytes and dendritic cells.

### 2.5 Sample Preparation for Glioblastoma scRNA-seq Data

The establishment and characterization of glioblastoma stem-like cells (GSCs) have been previously reported [49]. Briefly, GSCs were cultured in Dulbecco’s modified Eagle’s medium (DMEM)/F12 (Life Technologies) containing a B27 supplement minus vitamin A (Life Technologies), epidermal growth factor, and fibroblast growth factor 2 (20 ng/ml each; Wako Pure Chemicals Industries). For in vitro differentiation, GSCs were cultured in Dulbecco’s modified Eagle’s medium/F-12 medium (Life Technologies) containing 10% foetal bovine serum for the indicated times. Single-cell suspensions of GSCs or serum-induced differentiated GSCs were subjected to droplet-based scRNA-seq library preparation with the Chromium Single Cell 3’ Reagent Kit v2 (10x Genomics), aiming for an estimated 2,000 cells per library and following the manufacturer’s instructions. The libraries were checked with a BioAnalyzer High Sensitivity Chip (Agilent), quantified with a KAPA Library Quantification Kit (Roche), and then sequenced on the Illumina HiSeq 2500 platform in rapid mode.

### 2.6 Performance Evaluation

We evaluated the effectiveness of EEI compared to four existing methods: the Pearson correlation coefficient (referred to as Pearson), minet [50], GENIE3 [51] and PIDC [17].

The Pearson correlation is a basic correlation measure for a linear relationship between two variables ranging from -1.0 to 1.0. Note that the relationship of mutual exclusivity indicates that the Pearson coefficient is negative and greater than −1.0. minet and PIDC are methods for inferring of GRNs based on mutual information. minet infers the nonlinear relationship between two genes from microarray data and ranges 0.0 to 1.0. PIDC is an inference algorithm used to quantify the statistical relationships between triplets of genes based on the conditional mutual information from scRNA-seq data as positive values. GENIE3 infers the non-linear interactions among two genes based on a random forest regression and ranges 0.0 to 1.0. Since a very large amount of computational time is required for the large-scale datasets, we performed parallel computing by using 25 cores.

### 2.7 Performance Metrices

We evaluated the performance of EEI on the basis of the area under the precision-recall curve (AUPR) and average precision. The PR curve is plotted as the precision against recall. It is appropriate for binary classification with imbalanced data in which the number of positive samples is lower than the number of negative samples because the PR curve is sensitive to class distribution. Average precision (AP) indicates the weighted mean of precisions, with an increase in recall at each threshold (see Supplementary Appendix).

To validate the performance for clustering of single cells, we adopted the silhouette coefficient, which measures the cluster cohesion and separation and ranges −1.0 to 1.0. If the distance between one cluster and the other cluster is large, the silhouette coefficient is high. When evaluating the community detection of coexpression networks, we used the Szymkiewicz-Simpson coefficient and Jaccard index to measure the similarity between two sets of nodes, which ranged 0 to 1.

## 3 Results

### 3.1 Comparison of Exclusively Expressed Genes

#### 3.1.1 Comparison of Mutually Exclusive Gene Sets

Initially, we evaluated the effectiveness of EEI for the detection of mutually exclusive gene pairs by comparing it with four existing methods, the Pearson correlation coefficient, minet, GENIE3 and PIDC, using glioblastoma stem-like cell scRNA-seq data. Since minet, GENIE3 and PIDC cannot be applied to large-scale datasets, we used the expression data of the top 5,000 highly variable genes (HVG dataset) detected by Seurat [52]. Gene sets with mutually exclusive expression may exhibit genetic alterations and negative correlations. Highly mutually exclusive means that there is a high possibility of exclusive expression between two genes, rather than being a coincidence.

Since the five methods commonly output the prediction score for each gene pair, we assessed the performance of these methods for a binary classification problem. In this classification, we prepared 17 mutually exclusive gene pairs that are reported in the literature as positive samples (Table S1). We also prepared 50 negative gene pairs that were randomly sampled from 5,000 highly variable genes. To evaluate the performance of these methods, we used AUPR and AP scores (see Methods).

Table 1 and Figure S1(A) summarize the performances of the five methods. The AUPR and AP of EEI were the highest among all methods. While EEI could capture mutually exclusive patterns according to binary quantification, minet, GENIE3 and PIDC exhibited lower AUPR values, indicating that these methods could not correctly capture the exclusivity of expression due to excessive zero read counts.

**Table 1:** The prediction accuracies of five methods.

|  | EEI | Pearson | minet | GENIE3 | PIDC |
| --- | --- | --- | --- | --- | --- |
| AUPR | 0.51 | 0.47 | 0.45 | 0.40 | 0.42 |
| AP | 0.52 | 0.48 | 0.43 | 0.42 | 0.44 |

As shown in Figure S1(B), there were a few gene pairs had high EEI scores, while many other gene pairs presented lower EEI scores. Since 6 positive gene pairs were among the top pairs, actual mutually exclusive gene pairs could be predicted with moderate and high scores could be predicted by using EEI. Notably, the *PDGFRA* and *MET* gene pair showed the highest EEI of 12.4. Although minet and GENIE3 produced the highest scores for that pair, these methods inferred lower scores for other gene pairs. Therefore, these results indicate that our method enables us to identify mutually exclusive gene sets independent of the sequencing depth in sparse scRNA-seq data.

#### 3.1.2 Robustness Analysis of Read Depth in scRNA-seq Data

Since EEI identifies mutually exclusive gene pairs without taking into account technical zeros by dropouts, we examined the robustness of EEI against an insufficient read depth by comparison with the four methods. (Note that since PIDC cannot read the expression files, we used the other four methods.) We generated synthetic datasets from glioblastoma scRNA-seq data by randomly decreasing the total number of read counts from the original data by 10%.

First, we analysed the robustness for common gene pairs using the HVG dataset. We calculated the Pearson coefficient of the expression of two genes detected by minet and GENIE3 and regarded the top 500 gene pairs with negative coefficients as mutually exclusive gene pairs. Since the four methods shared 270 common gene pairs among the top 500 mutually exclusive gene pairs, we prepared these gene pairs as the positive samples and 500 negative samples that were randomly generated from genes that did not have zero expression. Figure 2 and Table S2 summarize the performances of the four methods, and EEI showed the best performance among them. Existing methods showed lower accuracy due to insufficient read counts as the read depth decreased. In contrast, EEI exhibited better performance at a lower sequencing read depth and was not strongly affected by a decrease in the total number of read counts.

**Figure 2:**
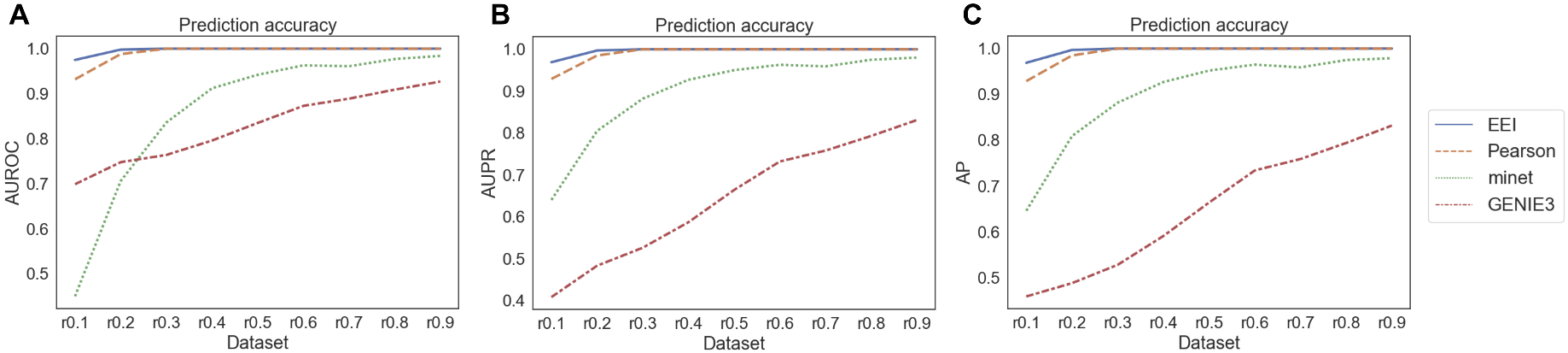
Comparison of the performances of the four methods with common gene sets from the glioblastoma scRNA-seq dataset. r0.1 represents the synthetic dataset in which 90% of read counts present zero expression compared to the original data. The average AUROC (**A**), AUPR (**B**), and AP (**C**) were calculated by repeating each simulation 10 times.

Second, we also analysed the robustness for gold standard gene pairs using two types of datasets: an HVG dataset and an expression dataset consisting of 18,597 genes that are expressed in at least one cell (NTZ dataset). We prepared 17 and 29 positive samples reported in the literature (listed in Table S1) for the HVG dataset and NTZ dataset, respectively and 50 negative samples. Since minet and GENIE3 cannot be applied to the large-scale datasets, we examined the performances of the EEI and Pearson methods. As the gene pairs of negative samples may also correspond to mutually exclusive gene pairs in some cases, we calculated only the AUPR and AP for the evaluation. As shown in Figure 3, Tables S3 and S4, EEI exhibited the best performance among all methods in both the HVG and NTZ datasets. For the HVG dataset, the AUPR and AP of Pearson, minet and GENIE3 were low and decreased as the lower sequencing depth decreased. In contrast, even when the sequence read depth decreased to 90% (r0.1 in Figure 3, Tables S3 and S4), the AUPR and AP of EEI showed moderate values.

**Figure 3:**
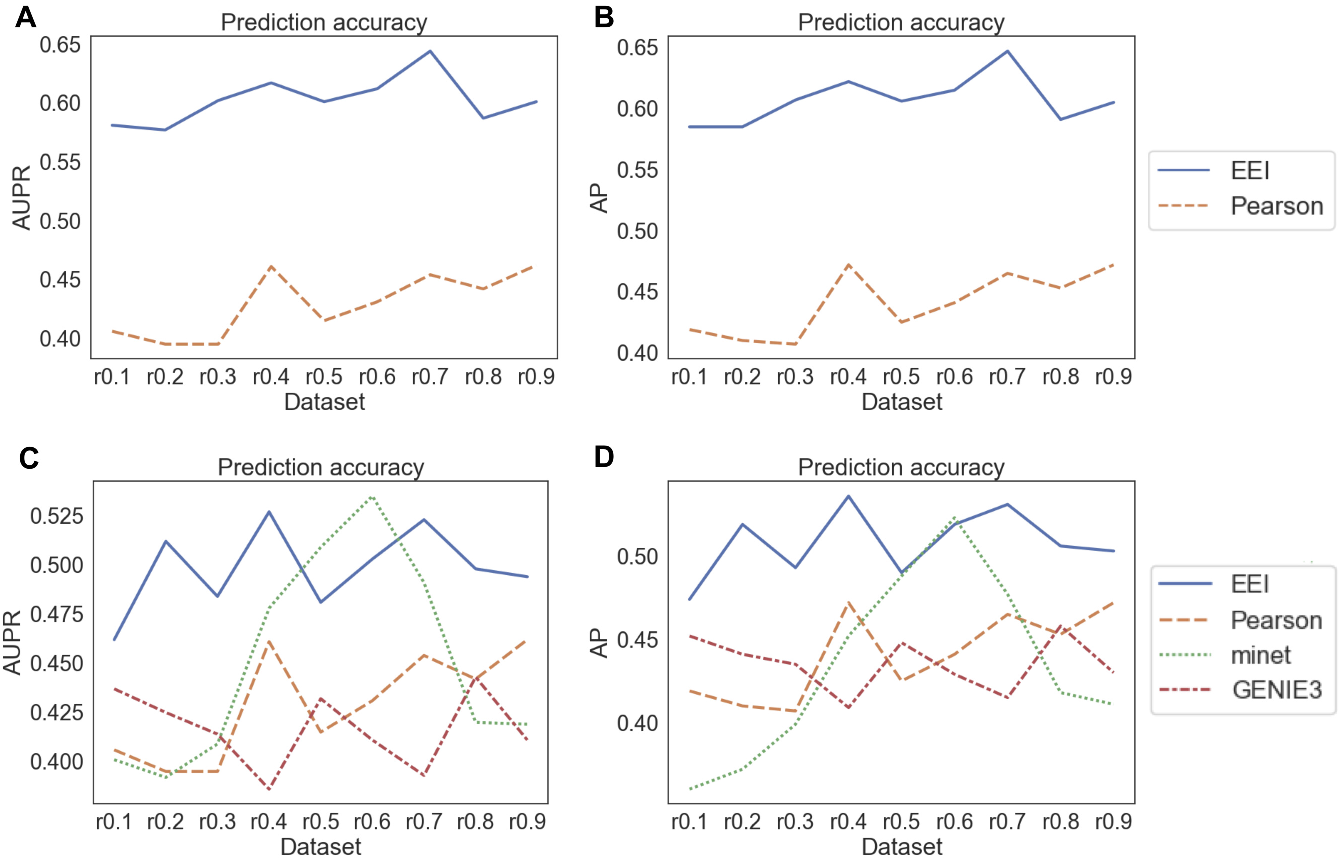
Comparison of the performances of the four methods with gold standard gene pairs. The AUPR (**A**) and AP (**B**) were calculated using the NTZ dataset and the AUPR (**C**) and AP (**D**) were calculated using the HVG dataset.

In particular, since the NTZ dataset contains a various types of genes, including not only highly variable genes but also genes with low expression, EEI could comprehensively capture various genes that were specifically expressed in each cell type. EEI enables the application of a large-scale dataset and the extensive capture of mutually exclusive expression levels even if a dataset is sparse. While the other methods were sensitive to the depth of sequencing, EEI was not affected by a decrease in the sequencing depth. These results suggest that our method enables us to comprehensively detect mutually exclusive gene sets while maintaining robustness against the sequencing read depth in sparse scRNA-seq data.

#### 3.1.3 Identification of Cell Marker Genes

To examine the possibility of the identification of marker genes, we assessed the performance of EEI for detecting marker genes by comparing the marker selection method and the databases for single cells. SCMarker [53] is an unsupervised marker selection method that identifies genes that are discriminatively expressed across cell types based on a mixture distribution model and are co- or mutually exclusively expressed. We calculated the prediction accuracy of EEI and SCMarker in terms of the positive and negative samples in Table S1(c) using human ES cells [44] and glioblastoma scRNA-seq datasets. The marker genes of human ES and cancer stem cells in glioblastoma have been reported in the previous studies [44] and [54, 55, 56], respectively. Note that since the two methods detected the different numbers of genes, we generated the same number of false positives and true negatives with SCMarker by randomly selecting genes detected from EEI.

For the glioblastoma dataset, EEI identified 5,354 genes above the threshold, 1.0, and SCMarker identified 456 genes. Table 2 shows that EEI outperformed SCMarker for detecting marker genes from the datasets that contain both sparse and sufficient read counts. We also analysed the detected gene pairs in glioblastoma in two public databases for cell type markers, CellMarker [57] and PanglaoDB [58], by EEI. The results showed that *PDGFRA* was included in CellMarker and that *PDGFRA, MET, MEF2C, OLIG1, SDC2, A2M, CHL1, MEG3* and *SLC1A3* were included in PanglaoDB. These markers are expressed in specific cell types in brain tissue. These results suggest that EEI has the possibility of detecting not only mutually exclusive gene pairs but also cell type marker genes.

**Table 2:** Prediction accuracies of EEI and SCMarker.

(a) Human ES dataset. (b) Glioblastoma dataset.
|  | EEI | SCMarker |  | EEI | SCMarker |
| --- | --- | --- | --- | --- | --- |
| Accuracy | 1.00 | 0.824 | Accuracy | 0.687 | 0.682 |
| Precision | 1.00 | 0.00284 | Precision | 0.0194 | 0.00439 |
| Sensitivity | 0.750 | 0.313 | Sensitivity | 0.643 | 0.143 |
| F1 score | 0.857 | 0.00564 | F1 score | 0.0377 | 0.00851 |

#### 3.1.4 Application of EEI to Single-Cell Clustering

In the classification of single cells, feature selection is an important step. Notably, it is crucial for there to be an association between gene expression features extracted from scRNA-seq data and the clustering of single cells. At the single-cell level, mutually exclusive gene sets due to genetic alterations might be considered to be expressed exclusively in different types of cells, which leads to tumour heterogeneity in cancer progression. This means that mutually exclusive gene sets can be used as features for the clustering of single cells. To evaluate the effectiveness of EEI, we compared the performances of the five methods using the five scRNA-seq datasets, as shown in Section 2.4. The feature matrix was generated by merging the ratio matrix of the top 1,000 mutually exclusive gene pairs by each method listed in Table S5 and the expression matrix of highly variable genes.

Figure 4 shows the UMAP results with EEI, the Pearson correlation, minet, GENIE3 and PIDC. For both the human ESC-derived neuron [46] and PBMC_CELseq2 [48] datasets, clustering with EEI showed that each cell type was detected as a separate cluster and produced compact clusters for all cell types. As shown in Figures S3 and S5, the silhouette coefficients of EEI were greater than those of the other methods for both datasets. For the other datasets shown in Figures S2, S4 and S6, the silhouette coefficients of PIDC and minet were higher than those of EEI for the human ES [44], mouse brain [47] and PBMC_MARSseq [48] datasets, respectively. In particular, since the PBMC_CELseq2, PBMC_MARSseq and human ES datasets were generated by CELseq2, MARSseq and Smartseq2 protocols, not by a droplet-based protocol, they contained sufficient read counts. In most cases, clustering with EEI displayed distinct clusters of cell types in these datasets. This means that EEI tends to be effective not only for sparse scRNA-seq data but also for data with a sufficient sequencing depth. Therefore, EEI enables us to capture the exclusive expression between two genes displaying intercell-type (but not intracell-type) heterogeneity.

**Figure 4:**
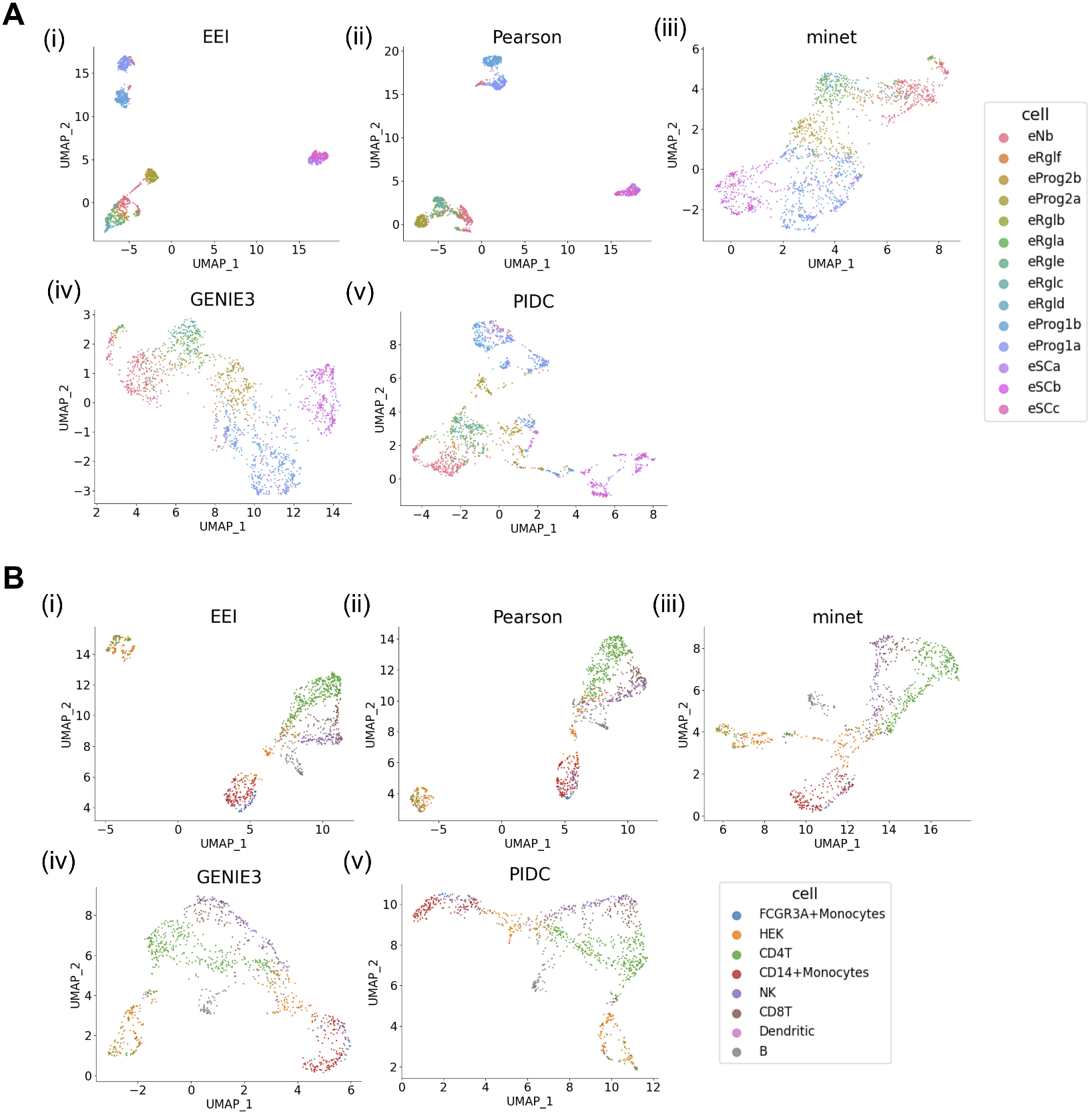
Comparison of the EEI (i), Pearson correlation (ii), minet (iii), GENIE3 (iv) and PIDC (v) UMAP results using human ES cell (**A**) and PBMC_CELseq2 (**B**) datasets.

In addition, we examined the effectiveness of EEI using human MEP scRNA-seq data [59]. This dataset consists of normalized data, including negative values, and does not contain any zero values. While CDI identified 854 gene pairs under a threshold of 0.01, EEI outputted no gene pairs. One reason for this result is that since the dataset does not include any zero values, EEI could not capture exclusive patterns by counting only the number of cells according to binary quantification. Therefore, these results suggest that EEI is widely applicable to various scRNA-seq datasets generated from distinct platforms and that the mutually exclusive gene sets detected by EEI can be applied to improve the sensitivity of the identification of cellular heterogeneity.

### 3.2 Comparison of Community Detection

#### 3.2.1 Community Detection using Human ES Cell scRNA-seq Data

First, we evaluated the performance of community detection with CDI for human embryonic stem cell scRNA-seq data [44] with a sufficient sequencing depth. For the differentiation of ES cells into specific cell types, several marker genes that are expressed in each cell type as reported in [44] were examined. The coexpression network was constructed under a CDI threshold of 10.0. A total of 3,270 genes were included in this network, and we identified 102 communities in total, including small-sized communities. Since the modularity score was 0.64, the network contained densely connected communities. Regarding known marker genes, we identified 13 markers described in [44] and analysed which markers were included in individual communities. Table 3 shows the marker genes and the cell types in which the corresponding markers were expressed.

**Table 3:** List of human ES cell type-specific genes included in each community and the corresponding cell types. Six genes specific to DE cells are included in the c0 community.

| Cell type name | Marker gene (Community No.) |
| --- | --- |
| H1 cells | *DNMT3B*(c97) |
| NPCs | *SOX2*(c97), *PAX6*(c0), *MAP2*(c97) |
| ECs | *CD34*(c98) |
| TBs | *GATA3*(c98), *HAND1*(c99) |
| DECs | *CER1*(c0), *EOMES*(c0), *GATA6*(c0), *LEFTY1*(c0),<br>*SOX17*(c0), *CXCR4*(c0) |

The detected marker genes were mostly included only in communities c0, c97, c98 and c99. In particular, 6 genes (*CER1, EOMES, GATA6, LEFTY1, SOX17* and *CXCR4*) specific to definitive endoderm (DE) cells were identified in the c0 community. Since human ES cells can differentiate towards DE cells, it is possible that c0 contains not only marker genes but also some genes associated with endoderm development. Similarly, other markers were mainly included in the c97 community, and it is possible that c97 contains sets of genes that exhibit functions specific to H1 and NP cells. Therefore, community detection enables us to identify groups of functionally related genes, in contrast to the analysis for individual genes using a coexpression network. These results suggest that CDI can capture coexpression patterns not only from sparse datasets but also from different types of datasets with a sufficient read depth.

#### 3.2.2 Comparison of Coexpressed Gene Sets

Second, we evaluated the performance of the community detection of gene coexpression networks from sparse scRNA-seq data. Since CDI can be applicable to large-scale networks, we focused on detecting communities of coexpression networks using the Leading eigenvector method (see Methods). We compared codependent gene sets by CDI using glioblastoma 0h scRNA-seq data with those by the Pearson correlation and cosine similarity. In this experiment, we selected the top 11,100 gene pairs under a CDI threshold of 10.0, and the same number of gene sets with positive coefficients in descending order were subjected to the other methods. The numbers of genes included in the resulting networks associated with CDI, the Pearson correlation and cosine similarity were 835, 1,221 and 436, respectively. The modularity score and the number of communities detected under each method are summarized in Table 4.

**Table 4:** Comparison of modularity scores and the numbers of communities under three methods applied to glioblastoma scRNA-seq data.

|  | CDI | Pearson correlation | Cosine similarity |
| --- | --- | --- | --- |
| Modularity | 0.38 | 0.35 | 0.09 |
| The number of communities | 8 | 147 | 29 |

The highest modularity score was 0.38 for CDI, and it was observed that the coexpression network according to CDI was divided into 6 medium-sized communities composed of approximately 10–300 genes (c0, c3, c4, c5, c6 and c7), as shown in Figure 5. In particular, the 0h_c3 community consisted of many genes that exhibited a high degree and were densely connected. At 0h, c0, c4 and c5 contained fewer hub genes than other communities. In contrast, the network obtained by the Pearson correlation was divided into several medium-sized and many small communities. Networks with high modularity scores presented sets of nodes that were densely connected.

**Figure 5:**
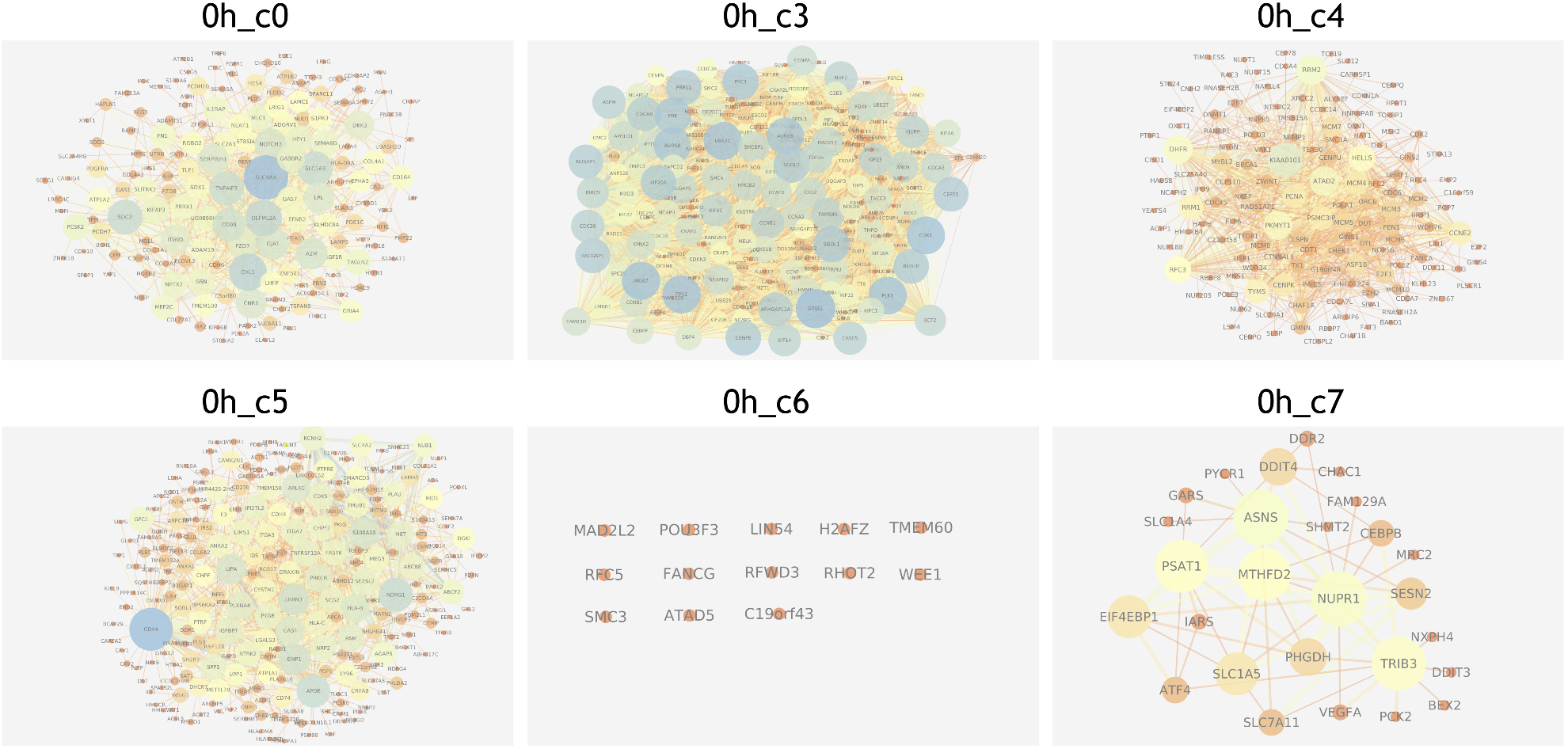
The structures of 6 extracted communities of coexpression networks by using CDI in the glioblastoma dataset. The blue node has a high degree. The 0h_c3 community contains many hub genes that are connected to many other genes.

To evaluate the properties of the communities obtained with the three methods, we cal-culated the Jaccard index to measure the similarity of the sets of genes in the community under each method. We regarded 30 genes included in the glioma pathway (hsa05214) in the Kyoto Encyclopedia of Genes and Genomes (KEGG) [60] as the gold standard: EGF, *TGFA, PDGF, IGF1, EGFR, PDGFRA, IGF1R, PLCG1, SHC1, GRB2, CALM, PRKCA, SOS, CAMK1, HRAS, PIK3CA, PTEN, BRAF, AKT, MAP2K1, MTDR, ERK, MDM2, TP53, P21, P16, CCND1, CDK4, RB1* and *E2F1.* Since there were some variations in community size, we used only communities containing more than 8 genes. The largest coefficients were 0.0156 for c0 and 0.0115 for c4. On the other hand, no genes identified according to the Pearson correlation and cosine similarity were shared with the KEGG pathways. The c0 community consisted of 165 genes, including the stemness marker genes *PDGFRA, A2M* and *NEAT1,* and the genes shared with the KEGG pathways were *PDGFRA, IGF1R* and *EGFR.* Interestingly, these genes were commonly included in the c0 community. We also performed Gene Ontology (GO) term enrichment analysis [61, 62], and 24 functional pathways were found to be significantly enriched (with a *p-value* of less than 1e — 04 for three shared genes), as listed in Figure S7. The top 3 pathways were *transmembrane receptor protein tyrosine kinase activity, positive regulation of DNA replication* and *tyrosine-protein kinase, catalytic domain.* Although there were a few shared genes between the 0h c0 community and the KEGG pathways, many different pathways were enriched compared to other CDI communities. There is a possibility that several specific genes associated with glioblastoma and various biological functions were included in the 0h c0 community. CDI can capture coexpression patterns from sparse scRNA-seq data if the sample size is sufficient. However, it must be noted that it does not necessarily provide information indicating that two genes are positively correlated. These results suggest that the coexpression network associated with CDI contained densely connected subnetworks and that community detection enables us to identify not only possible densely connected subgraphs but also biologically functional networks.

#### 3.2.3 Comparison of Coexpression Networks for Multiple Samples

To validate the effectiveness of community detection when multiple samples are provided, we performed a comparative analysis of coexpression networks from glioblastoma scRNA-seq data at different time points. The dataset for 0 hours consisted of 18,597 genes and 2,102 cells, and the dataset from 12 hours consisted of 18,163 genes and 2,209 cells. Using these datasets, we reconstructed coexpression networks based on a CDI threshold of 10.0. After decomposition of the networks using the Leading eigenvector method, to compare extracted communities, we calculated the Szymkiewicz-Simpson coefficient to evaluate the similarity between two communities at different time points.

As shown in Figure 6, every network exhibited a power-law degree distribution. The scale-free network presented an uneven node degree distribution and contained a few hub nodes. With regard to the hub genes, in the 0-hour network, *CDK1, PBK, KIAA0101, GTSE1, MKI67, UBE2C, SGOL1, TPX2, AURKB* and *RRM2* were identified, and in the 12-hour network, *TOP2A, PBK, TYMS, NUSAP1, BIRC5, UBE2C, TPX2, GTSE1, ATAD2* and *MKI67* were also identified. Additionally, the results showed that 4 communities including more than 100 genes were extracted for each sample. We observed high similarity for shared communities: 0.83 for 0h c3 and 12h_c6 and 0.74 for 0h_c4 and 12h_c7 (see Table S6). In contrast, 0h c0 and 12h_c8 as well as 0h c5 and 12h_c0 were considered to be specific communities in response to external stimuli. For example, it has been reported that the known stemness marker genes *PDGFRA* and *MET* exhibit different expression patterns [34]. While *PDGFRA* and neighbouring genes existed in 0h c0, *PDGFRA* did not exist in any community, and some neighbouring genes remained in 12h_c8. *MET* and neighbouring genes existed in both in 0h c5 and 12h c0. EEI results showed that *PDGFRA* and *MET* were expressed in a mutually exclusive manner, indicating that these two genes are expressed in the different cell populations [34]. This means that although there are some variations in the expression of individual genes, the sets of genes that constitute the community are partially conserved regardless of the effects of external stimuli. Therefore, these results suggest that community-based comparisons of coexpression networks enable us to detect similarities and differences in coexpressed genes across multiple samples.

**Figure 6:**
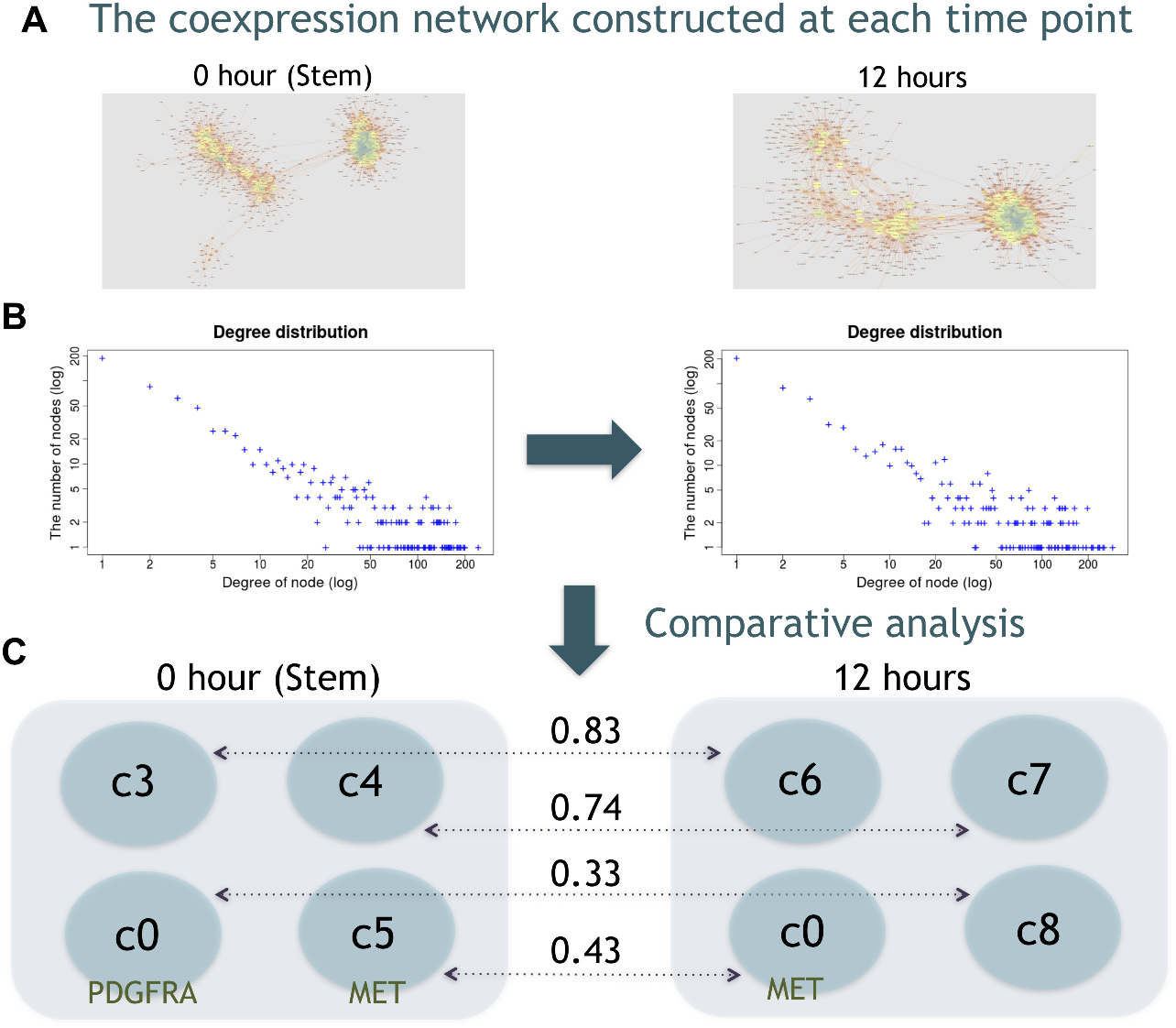
Comparative analysis of coexpression networks at different time points. The coexpression network constructed at each time point (**A**) and degree distribution (**B**). The blue and yellow nodes represent the high- and low-degree nodes, respectively. After decomposition of the coexpression networks, four communities were extracted for each sample, and the values between the communities at 0 and 12 hours represent the Szymkiewicz-Simpson coefficients (**C**).

### 4 Conclusion and Discussion

In this study, we developed a novel metric, EEI, to comprehensively quantify mutual exclusivity between two genes from sparse scRNA-seq data. A comparison with existing methods using glioblastoma scRNA-seq data suggested that EEI identified gene sets due to genetic alterations and negative correlations.

In particular, our findings show that EEI is effective for detecting mutually exclusive gene sets, while maintaining robustness against the sequencing read depth in droplet-based scRNA-seq data. We also applied EEI to improve the sensitivity of the classification of single cells. The results suggest that exclusive expression can be introduced to identify intercell-type heterogeneity based on the feature matrix.

We also examined the performance of coexpression networks from glioblsatoma scRNA-seq data in community detection. Although the Louvain method is faster, we used the Leading eigenvector method, which is fundamental and sufficiently applicable to large-scale networks. The results suggested that the communities detected from CDI contained more densely connected subgraphs than existing methods, and some marker genes associated with specific pathways in glioma were identified. Community detection enables us to identify candidate marker genes from known marker genes. A community-based comparison provides information not only on functionally related genes but also on the similarities and differences in coexpressed genes when multiple samples are provided.

Since EEI does not impute technical zeros and captures genes that are mutually exclusively expressed without discriminating those with zero expression due to biological and technical zeros, the imputed data might be able to improve the detection of gene pairs. Although EEI and CDI can be applied to large-scale datasets, they require considerable computational time and are effectively parallelizable. In addition, while the mutually exclusive gene sets identified by EEI can improve the sensitivity of the identification of cell-to-cell heterogeneity, this approach is not suitable for datasets containing excessive zeros, and another future goal will be to improve the extraction of expression features.

### 5 Data Availability

The code to calculate CDI and EEI is available at https://github.com/Natsu01/EEISP. The raw sequencing data and processed files of the glioblastoma scRNA-seq are available at the NCBI Gene Expression Omnibus (GEO) under accession number GSE144623.

## 6 Acknowledgements

This research was partially supported by Platform Project for Supporting Drug Discovery and Life Science Research (Basis for Supporting Innovative Drug Discovery and Life Science Research (BINDS)) from AMED under grant number JP19am0101105.

## 8 Author Contributions

N.N. performed the implementation and computational studies: T. Akutsu, and R.N. organized the project: T.H. and T. Akiyama prepared the glioblastoma scRNA-seq data: K.F. and K.S. sequenced the scRNA-seq data: N.N. and R.N. wrote the draft of the manuscript, and all authors approved the submitted version.

