## Supplementary Material for "Codependency and mutual exclusivity for gene community detection from sparse single-cell transcriptome data"

---

### SUPPLEMENTARY MATERIAL

### 1 Supplementary Appendix

#### 1.1 Performance metrics

To evaluate the performance of EEI for detecting mutually exclusive gene sets, we calculated the prediction accuracy, AUROC, AUPR and average precision (AP). The AUROC is plotted as the true positive rate (TPR) against the false positive rate (FPR). The AUPR is plotted as the precision against recall. The average precision (AP) indicates the weighted mean of precisions with an increase in recall at each threshold. These metrics are defined as follows.

$$\begin{aligned} TPR(recall) &= \frac{TP}{TP+FN}, \\ FPR &= \frac{FP}{FP+TN}, \\ Precision &= \frac{TP}{TP+FP}, \\ AP &= \sum_n (R_n - R_{n-1})P_n, \end{aligned} \tag{1}$$

where TP, FP, TN and FN indicate the numbers of true positives, false positives, true negatives and false negatives, respectively, and $R_n$ and $P_n$ indicate the recall and precision at the $n$-th threshold, respectively.

In the performance evaluation of the community detection of coexpression networks, we used the Szymkiewicz-Simpson coefficient and Jaccard index to measure the similarity between two sets of nodes, ranging from 0 to 1, which were calculated as follows:

$$overlap(A,B) = \frac{|A \cap B|}{(\min |A|, |B|)}, \tag{2}$$

$$J(A,B) = \frac{|A \cap B|}{|A \cup B|}, \tag{3}$$

where $A$ and $B$ represent the sets of nodes.

### 2 Supplementary Figures

#### 2.1 Comparison of Mutually Exclusive Gene Sets

We evaluated the performances of the five methods using the glioblastoma dataset. Figure S1 shows the AUPR and the score distributions of EEI, the Pearson correlation, minet, GENIE3 and PIDC.

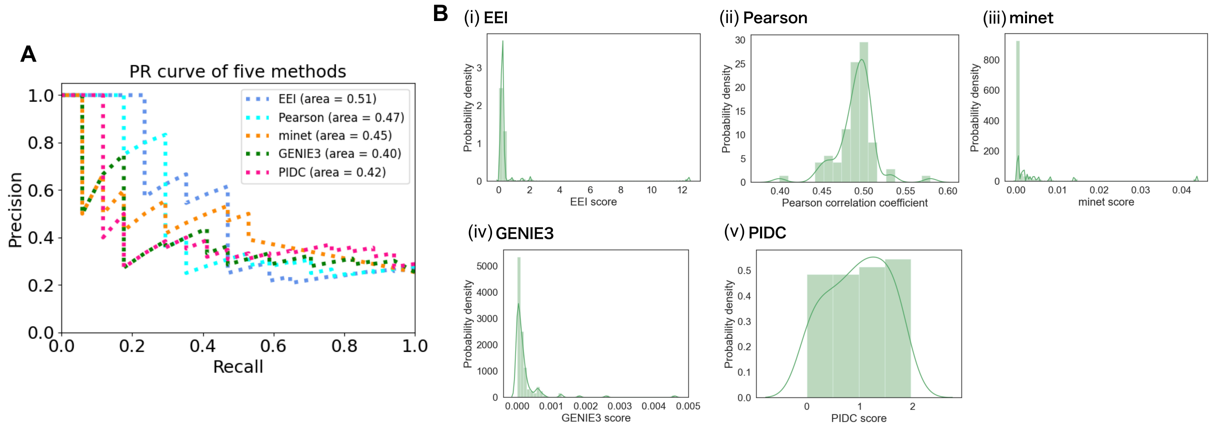

Supplementary Figure S1: The PR curves of the five methods for the glioblastoma dataset (**A**). Comparison of the distributions of the EEI score (i), the Pearson correlation coefficient (ii), the minet score (iii), the GENIE3 score (iv) and the PIDC score (v) (**B**). The x-axis, y-axis and curve represent the coefficient, probability density and probability density function, respectively.

#### 2.2 Application of EEI to Single-Cell Clustering

We tried to apply mutually exclusive gene sets to improve the sensitivity of the identification of single cells. To compare the performance, we applied the five methods to the five scRNA-seq datasets. When generating the feature matrix, we used 1,000 mutually exclusive gene sets for each of the five methods. Figures S2, S3, S4, S5 and S6 show the UMAP results of the five methods and the average silhouette coefficient for each clustering with human ES cells [1], human ESC-derived neurons [2], mouse brain cells [3], PBMC_CELseq2 cells [4] and PBMC_MARSseq cells [4], respectively.

#### 2.3 Comparison of Coexpressed Gene Sets

We performed Gene Ontology (GO) term enrichment analysis for three shared genes, *PDGFRA*, *IGF1R*, and *EGFR*, and 24 functional pathways with a *p-value* of less than $1e-04$ were annotated, as shown in Figure S7.

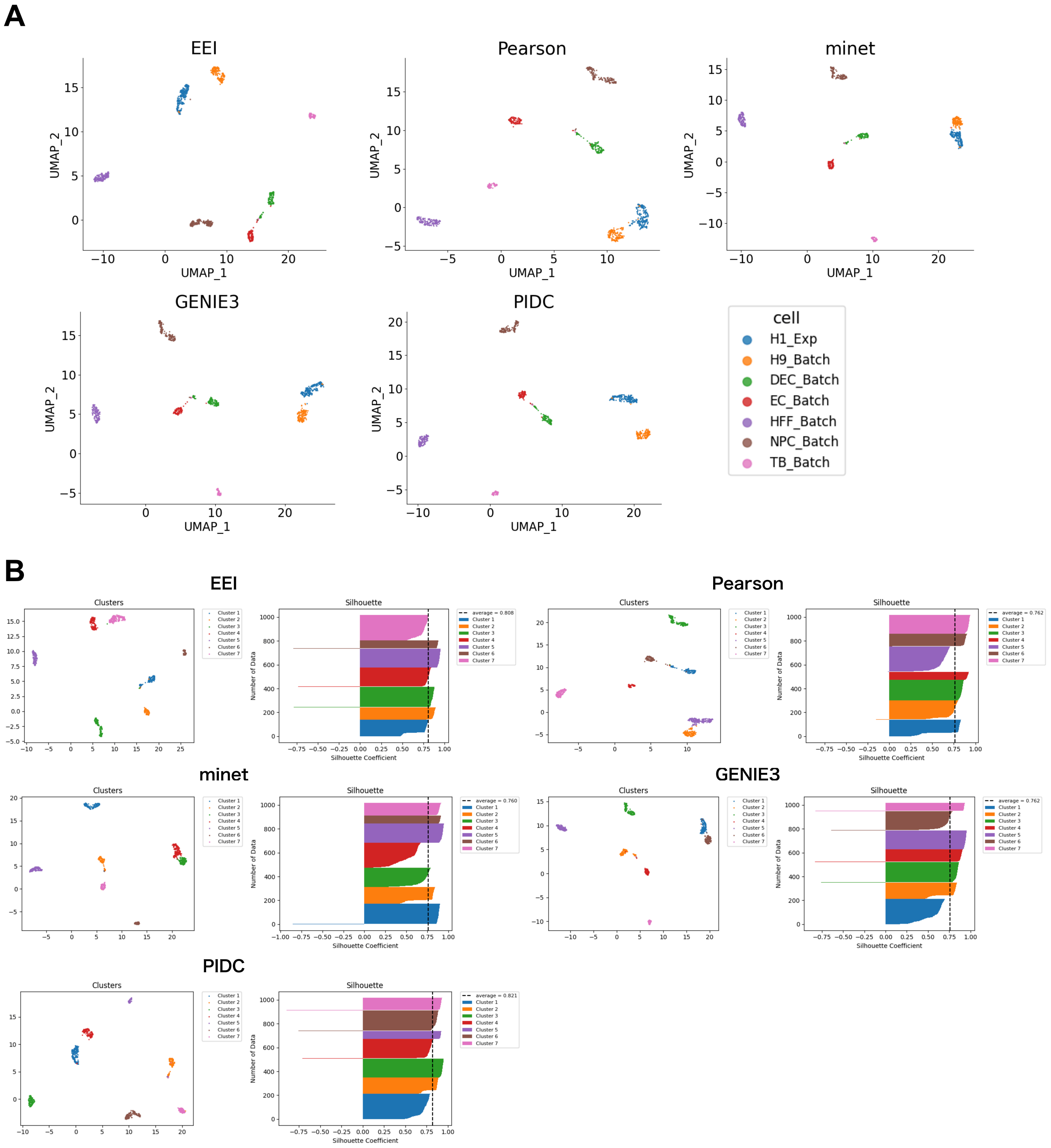

Supplementary Figure S2 : Comparison of the EEI, Pearson, minet, GENIE3 and PIDC UMAP results using scRNA-seq data from human ES cells (**A**). Comparison of the silhouette coefficients for the five methods (**B**).

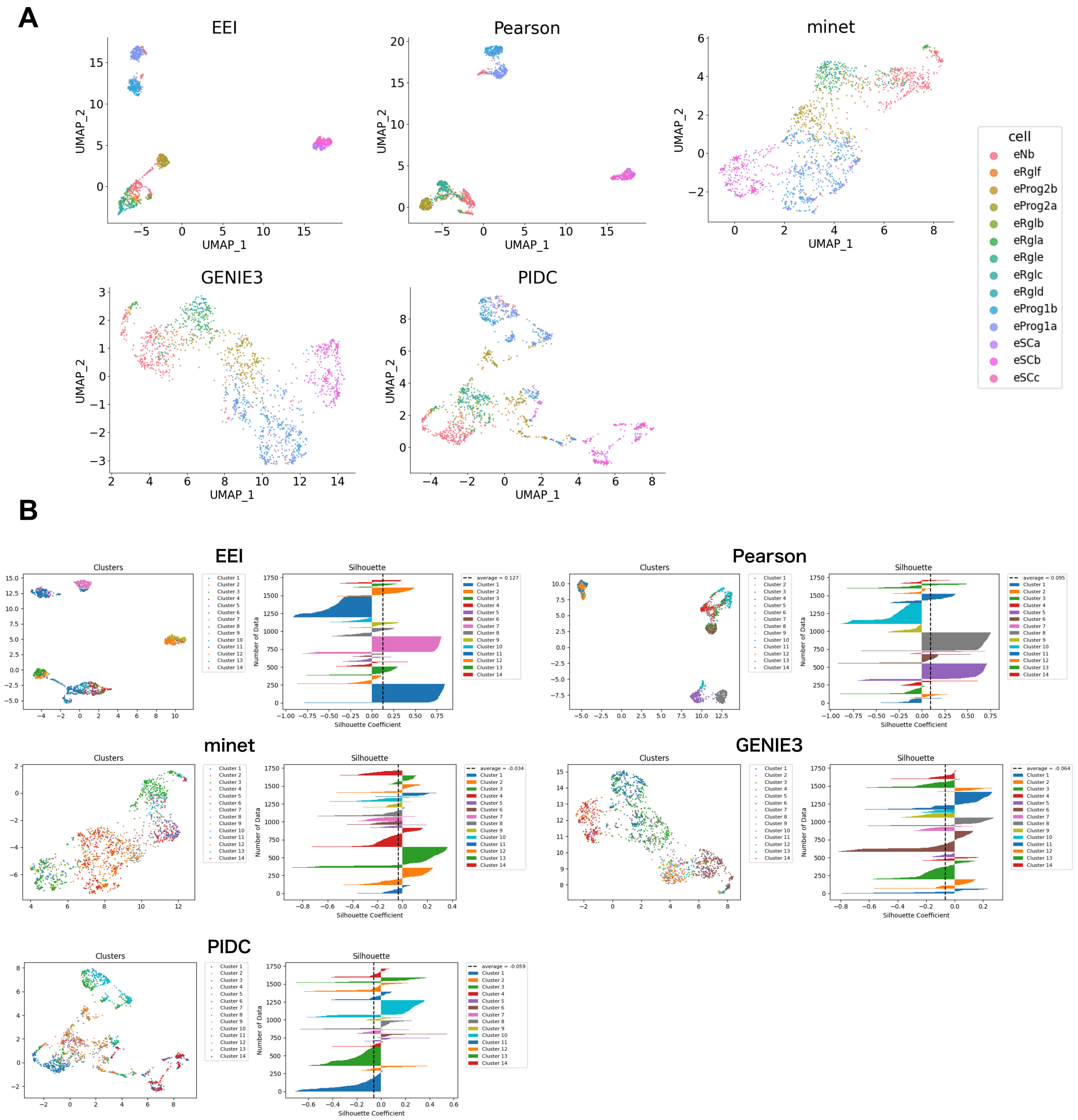

Supplementary Figure S3 : Comparison of the EEI, Pearson, minet, GENIE3 and PIDC UMAP results using scRNA-seq data from human ESC-derived neurons (**A**). Comparison of the silhouette coefficients for the five methods (**B**).

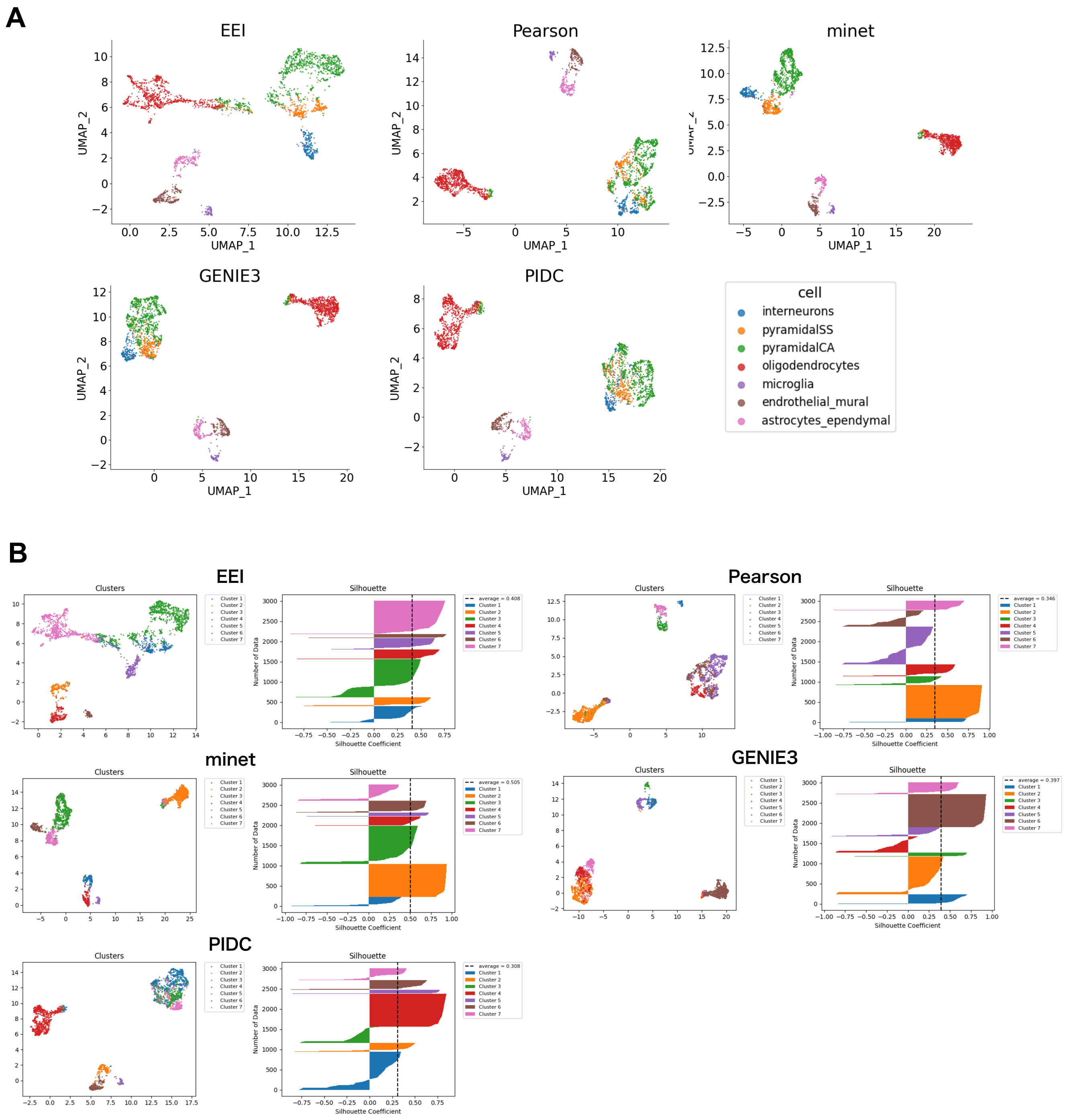

Supplementary Figure S4 : Comparison of the EEI, Pearson, minet, GENIE3 and PIDC UMAP results using scRNA-seq data from mouse brain cells (**A**). Comparison of the silhouette coefficients for the five methods (**B**).

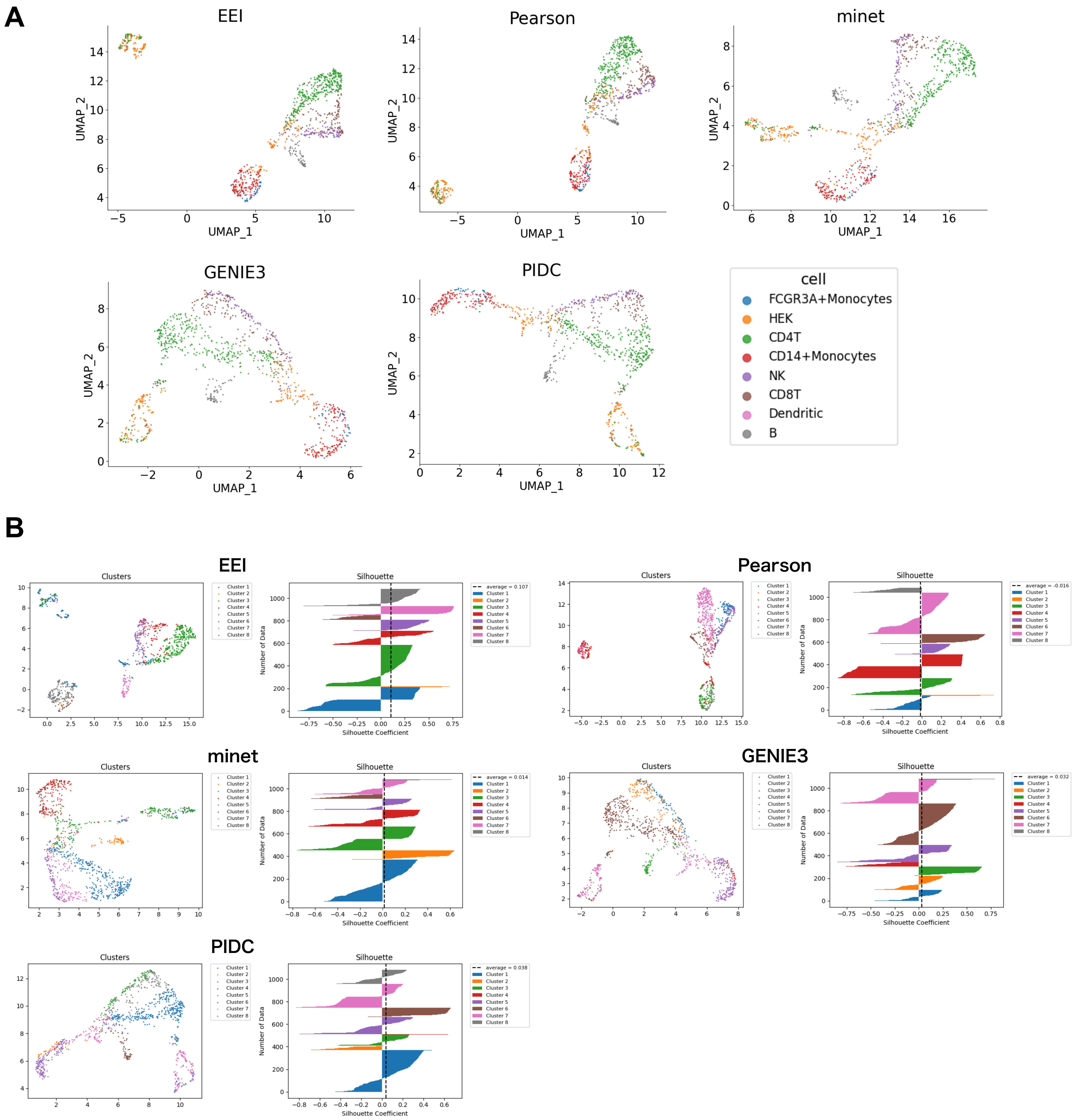

Supplementary Figure S5 : Comparison of the EEI, Pearson, minet, GENIE3 and PIDC UMAP results using scRNA-seq data generated by CELseq2 from PBMCs (**A**). Comparison of the silhouette coefficients for the five methods (**B**).

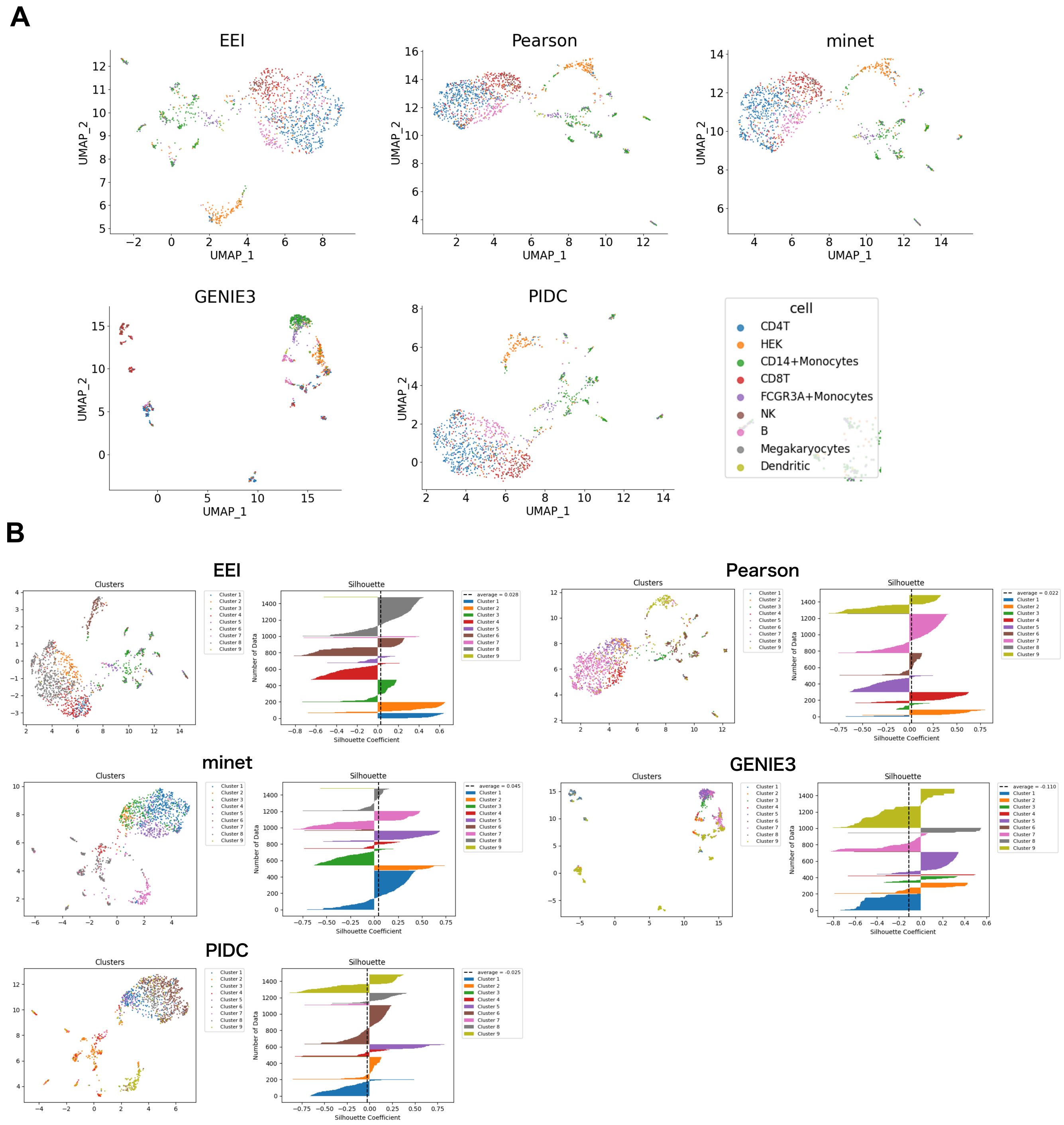

Supplementary Figure S6 : Comparison of the EEI, Pearson, minet, GENIE3 and PIDC UMAP results using scRNA-seq data generated by MARSseq from PBMCs (**A**). Comparison of the silhouette coefficients for the five methods (**B**).

| Category | Term | RT | Genes | Count | % | P-Value | Benjamini |
|---|---|---|---|---|---|---|---|
| GOTERM_MF_DIRECT | transmembrane receptor protein tyrosine kinase activity | RT | | 3 | 100.0 | 4.9E-6 | 1.9E-4 |
| GOTERM_BP_DIRECT | positive regulation of DNA replication | RT | | 3 | 100.0 | 6.1E-6 | 7.2E-4 |
| INTERPRO | Tyrosine-protein kinase, catalytic domain | RT | | 3 | 100.0 | 2.1E-5 | 3.0E-4 |
| INTERPRO | Tyrosine-protein kinase, active site | RT | | 3 | 100.0 | 2.6E-5 | 3.0E-4 |
| UP_KEYWORDS | Tyrosine-protein kinase | RT | | 3 | 100.0 | 2.9E-5 | 9.1E-4 |
| GOTERM_BP_DIRECT | phosphatidylinositol-mediated signaling | RT | | 3 | 100.0 | 3.9E-5 | 2.3E-3 |
| INTERPRO | Serine-threonine/tyrosine-protein kinase catalytic domain | RT | | 3 | 100.0 | 5.7E-5 | 4.3E-4 |
| SMART | TyrKc | RT | | 3 | 100.0 | 7.1E-5 | 4.2E-4 |
| KEGG_PATHWAY | Glioma | RT | | 3 | 100.0 | 8.8E-5 | 2.0E-3 |
| GOTERM_BP_DIRECT | protein autophosphorylation | RT | | 3 | 100.0 | 1.0E-4 | 3.5E-3 |
| KEGG_PATHWAY | Melanoma | RT | | 3 | 100.0 | 1.1E-4 | 2.0E-3 |
| GOTERM_BP_DIRECT | positive regulation of cell migration | RT | | 3 | 100.0 | 1.2E-4 | 3.5E-3 |
| KEGG_PATHWAY | Prostate cancer | RT | | 3 | 100.0 | 1.6E-4 | 2.1E-3 |
| BIOCARTA | Erk1/Erk2 Mapk Signaling pathway | RT | | 3 | 100.0 | 3.3E-4 | 8.2E-3 |
| INTERPRO | Protein kinase, ATP binding site | RT | | 3 | 100.0 | 4.2E-4 | 2.3E-3 |
| UP_SEQ_FEATURE | domain:Protein kinase | RT | | 3 | 100.0 | 5.7E-4 | 1.2E-2 |
| INTERPRO | Protein kinase, catalytic domain | RT | | 3 | 100.0 | 6.9E-4 | 2.3E-3 |
| INTERPRO | Furin-like cysteine-rich domain | RT | | 2 | 66.7 | 7.5E-4 | 2.3E-3 |
| INTERPRO | EGF receptor, L domain | RT | | 2 | 66.7 | 7.5E-4 | 2.3E-3 |
| GOTERM_BP_DIRECT | positive regulation of cell proliferation | RT | | 3 | 100.0 | 7.7E-4 | 1.8E-2 |
| UP_SEQ_FEATURE | binding site:ATP | RT | | 3 | 100.0 | 7.7E-4 | 1.2E-2 |
| INTERPRO | Protein kinase-like domain | RT | | 3 | 100.0 | 8.2E-4 | 2.3E-3 |
| KEGG_PATHWAY | Focal adhesion | RT | | 3 | 100.0 | 8.9E-4 | 6.8E-3 |
| KEGG_PATHWAY | Rap1 signaling pathway | RT | | 3 | 100.0 | 9.3E-4 | 6.8E-3 |

Supplementary Figure S7 : Twenty-four enriched pathways for three genes (*PDGFRA*, *IGF1R* and *EGFR*) shared with the KEGG pathways.

### 3 Supplementary Tables

#### 3.1 Comparison of Mutually Exclusive Gene Sets

We evaluated the performance of EEI for the detection of mutually exclusive gene pairs by comparing four existing methods, the Pearson correlation coefficient, minet, GENIE3 and PIDC using glioblastoma stem-like cell scRNA-seq data. We assessed the performances of the five methods for a binary classification problem and prepared 17 mutually exclusive gene pairs of the HVG dataset reported in the literature as positive samples listed in Table S1(a).

Supplementary Table S1: Mutually exclusive gene sets in positive samples in the performance evaluation.

(a) 17 gold standard gene pairs

| gene1 | gene2 | Ref. |
|---|---|---|
| PDGFRA | MET | [5] |
| EGFR | IGF2 | [6] |
| EGFR | OLIG1 | [7] |
| EGFR | PDGFRA | [5] |
| EGFR | MET | [9] |
| CD24 | GFAP | [10] |
| CD24 | APOE | [10] |
| CD24 | AQP4 | [10] |
| CD24 | CD44 | [10] |
| CD24 | CD9 | [10] |
| CD24 | VIM | [10] |
| DCX | GFAP | [10] |
| DCX | APOE | [10] |
| DCX | AQP4 | [10] |
| DCX | CD44 | [10] |
| DCX | CD9 | [10] |
| DCX | VIM | [10] |

(b) 29 gold standard gene pairs

| gene1 | gene2 | Ref. |
|---|---|---|
| PDGFRA | MET | [5] |
| PDGFRA | IDH2 | [11] |
| EGFR | IGF2 | [6] |
| EGFR | OLIG1 | [7] |
| EGFR | PDGFRA | [5] |
| EGFR | TP53 | [12] |
| EGFR | MET | [9] |
| NNMT | OLIG2 | [13] |
| KLF4 | FOXO1 | [14] |
| MDM2 | TP53 | [15] |
| ITGA6 | KLF9 | [16] |
| CD24 | GFAP | [10] |
| CD24 | APOE | [10] |
| CD24 | AQP4 | [10] |
| CD24 | CD44 | [10] |
| CD24 | CD9 | [10] |
| CD24 | VIM | [10] |
| DCX | GFAP | [10] |
| DCX | APOE | [10] |
| DCX | AQP4 | [10] |
| DCX | CD44 | [10] |
| DCX | CD9 | [10] |
| DCX | VIM | [10] |
| SOX11 | GFAP | [10] |
| SOX11 | APOE | [10] |
| SOX11 | AQP4 | [10] |
| SOX11 | CD44 | [10] |
| SOX11 | CD9 | [10] |
| SOX11 | VIM | [10] |

(c) 11 gold standard gene pairs

| gene1 | gene2 | Ref. |
|---|---|---|
| PDGFRA | MET | [5] |
| PDGFRA | IDH2 | [11] |
| EGFR | IGF2 | [6] |
| EGFR | OLIG1 | [7] |
| EGFR | PDGFRA | [5] |
| EGFR | TP53 | [12] |
| EGFR | MET | [9] |
| NNMT | OLIG2 | [13] |
| KLF4 | FOXO1 | [14] |
| MDM2 | TP53 | [15] |
| ITGA6 | KLF9 | [16] |

#### 3.2 Robustness Analysis of Read Depth in scRNA-seq Data

Since EEI identifies mutually exclusive gene sets without taking into account of imputing technical zeros, we performed a robustness analysis of the four methods against an insufficient read depth. Since minet and GENIE3 cannot be applied to large-scale networks, we evaluated the performances of the four methods using both the HVG and NTZ datasets. We regarded 29 mutually exclusive gold standard gene pairs as the positive samples (listed in Table S1(b)). Tables S2, S3 and S4 show the prediction accuracies of the four methods using the common gene pairs and gold standard gene pairs as the positive samples.

#### 3.3 Identification of Cell Marker Genes

We also evaluated the performances of the methods for detecting marker genes with 11 positive samples as shown in Table S1(c), by comparison with SCMarker [17].

#### 3.4 Application of EEI to the Clustering of Single Cells

In the clustering analysis, we used the top 1,000 gene pairs detected by EEI, Pearson, minet, GENIE3 and PIDC, and these are listed in Table S5.

Supplementary Table S2: Robustness analysis of the four methods using the common gene pairs. r0.1 represents that the synthetic dataset in which 90% zero expression values are included compared to the original data. The total read counts represent the average total number of reads included in the corresponding data.

| | | r0.9 | r0.8 | r0.7 | r0.6 | r0.5 | r0.4 | r0.3 | r0.2 | r0.1 |
|---|---|---|---|---|---|---|---|---|---|---|
| | The total read counts | 3272622 | 2909112 | 2544942 | 2181803 | 1818567 | 1454335 | 1091338 | 727300 | 353255 |
| EEI | AUROC | 0.96 | 1.00 | 1.00 | 1.00 | 1.00 | 1.00 | 1.00 | 1.00 | 1.00 |
| | AUPR | 0.97 | 1.00 | 1.00 | 1.00 | 1.00 | 1.00 | 1.00 | 1.00 | 1.00 |
| | AP | 0.97 | 1.00 | 1.00 | 1.00 | 1.00 | 1.00 | 1.00 | 1.00 | 1.00 |
| Pearson | AUROC | 0.93 | 0.99 | 1.00 | 1.00 | 1.00 | 1.00 | 1.00 | 1.00 | 1.00 |
| | AUPR | 0.93 | 0.99 | 1.00 | 1.00 | 1.00 | 1.00 | 1.00 | 1.00 | 1.00 |
| | AP | 0.93 | 0.99 | 1.00 | 1.00 | 1.00 | 1.00 | 1.00 | 1.00 | 1.00 |
| minet | AUROC | 0.45 | 0.71 | 0.84 | 0.91 | 0.94 | 0.97 | 0.96 | 0.98 | 0.98 |
| | AUPR | 0.64 | 0.81 | 0.88 | 0.93 | 0.95 | 0.97 | 0.96 | 0.98 | 0.98 |
| | AP | 0.65 | 0.81 | 0.88 | 0.93 | 0.95 | 0.97 | 0.96 | 0.98 | 0.98 |
| GENIE3 | AUROC | 0.70 | 0.75 | 0.76 | 0.80 | 0.84 | 0.87 | 0.89 | 0.91 | 0.93 |
| | AUPR | 0.41 | 0.48 | 0.53 | 0.59 | 0.66 | 0.73 | 0.76 | 0.79 | 0.83 |
| | AP | 0.46 | 0.49 | 0.53 | 0.59 | 0.66 | 0.73 | 0.76 | 0.79 | 0.83 |

Supplementary Table S3: Robustness analysis of EEI and Pearson with the gold standard gene pairs using the NTZ dataset.

| | | r0.9 | r0.8 | r0.7 | r0.6 | r0.5 | r0.4 | r0.3 | r0.2 | r0.1 |
|---|---|---|---|---|---|---|---|---|---|---|
| | The total read counts | 10584790 | 9408292 | 8231448 | 7056196 | 5880811 | 4703815 | 3527747 | 2351791 | 1176231 |
| EEI | AUPR | 0.58 | 0.58 | 0.60 | 0.62 | 0.60 | 0.61 | 0.64 | 0.59 | 0.60 |
| | AP | 0.97 | 1.00 | 1.00 | 1.00 | 1.00 | 1.00 | 1.00 | 1.00 | 1.00 |
| Pearson | AUPR | 0.41 | 0.40 | 0.40 | 0.46 | 0.42 | 0.43 | 0.45 | 0.44 | 0.46 |
| | AP | 0.42 | 0.41 | 0.41 | 0.47 | 0.43 | 0.44 | 0.47 | 0.45 | 0.47 |

Supplementary Table S4: Robustness analysis of the four methods using the gold standard gene pairs using the HVG dataset.

| | | r0.9 | r0.8 | r0.7 | r0.6 | r0.5 | r0.4 | r0.3 | r0.2 | r0.1 |
|---|---|---|---|---|---|---|---|---|---|---|
| | The total read counts | 3272622 | 2909112 | 2544942 | 2181803 | 1818567 | 1454335 | 1091338 | 727300 | 353255 |
| EEI | AUPR | 0.46 | 0.51 | 0.48 | 0.53 | 0.48 | 0.50 | 0.52 | 0.50 | 0.49 |
| | AP | 0.47 | 0.52 | 0.49 | 0.54 | 0.49 | 0.52 | 0.53 | 0.52 | 0.50 |
| Pearson | AUPR | 0.41 | 0.40 | 0.40 | 0.46 | 0.42 | 0.43 | 0.45 | 0.44 | 0.46 |
| | AP | 0.42 | 0.41 | 0.41 | 0.47 | 0.43 | 0.44 | 0.47 | 0.45 | 0.47 |
| minet | AUPR | 0.40 | 0.40 | 0.41 | 0.48 | 0.51 | 0.54 | 0.49 | 0.42 | 0.42 |
| | AP | 0.36 | 0.37 | 0.40 | 0.45 | 0.49 | 0.52 | 0.48 | 0.42 | 0.41 |
| GENIE3 | AUPR | 0.44 | 0.43 | 0.41 | 0.39 | 0.43 | 0.41 | 0.39 | 0.44 | 0.41 |
| | AP | 0.45 | 0.44 | 0.44 | 0.41 | 0.45 | 0.43 | 0.42 | 0.46 | 0.43 |
